# Spatial segregation and cross-kingdom interactions drive stingless bee hive microbiome assembly

**DOI:** 10.1101/2025.03.07.642116

**Authors:** Lílian Caesar, Carlin Barksdale, Victor Hugo Valiati, Irene Newton

## Abstract

Studying host-associated microbiome assembly is key to understanding microbial and host evolution and health. While honey bee microbiomes have been central models for such investigations among pollinators, they overlook the diversity of eusocial dynamics and multi-kingdom interactions. Stingless bees, highly eusocial managed bees that rely on yeast for larval development, offer a valuable complementary system to study microbiome assembly, and within an eco-evolutionary framework. Using amplicon sequencing, metagenomics, and microbial experiments, we investigate the drivers of stingless bee microbiome assembly. We reveal a spatially structured, site-adapted microbiome, where high microbial influx hive components are segregated from the brood, which harbors a stable, multi-kingdom community. We show that the brood microbiome is not only physically protected but also actively maintained through highly selective bacterial-fungal interactions. Our findings uncover multi-layered mechanisms shaping an eusocial insect microbiome, from host biology to cross-kingdom interactions, while providing critical insights into microbiome maintenance of important pollinators.

## Introduction

Microbiomes expand their host’s genetic repertoire, influencing digestion, immunity, behavior, and fitness^1–3^. Such benefits often rely on specific and balanced microbial communities, making microbiome assembly and maintenance crucial, and potentially driving host-microbiome codiversification throughout evolution^4–7^. Understanding the factors that shape microbiomes thus remains a central question in the field, with implications for uncovering the evolution of microbial and host features, predicting microbiome responses to environmental changes, and informing strategies for microbiome manipulation.

Multiple mechanisms shape microbiomes. For instance, host-to-microbe interactions influence community structure through gut compartmentalization^8^; microbe-to-host interactions secure key symbionts for diet digestion^9^; and microbe-to-microbe interactions support communities via metabolic cross-feeding^10^. The influence of mechanisms may vary across parts of an organism. For example, in plant microbiomes microbe-to-microbe interactions play important roles in rhizosphere and bulk soil microbiome assembly, while host-microbe interactions exert extra pressure in root and leaf endospheres^11,12^. In social organisms, host behavior may also influence microbial transmission, and body sites with high microbial influx, like oral microbiomes, may further impact microbiome assembly^13^. Expanding studies to diverse systems is thus key to uncovering the mechanisms of microbiome assembly.

Eusocial insects are powerful and complex systems for studying microbiome assembly due to their highly connected societies and protected colonies, where direct fitness is measured at the colony level^14^. The microbiome in each colony component, including individuals and hive sites like food stores, might be expected to be functionally integrated — as the microbiome of different body sites of an organism^11^. In honey bees (*Apis mellifera*), for example, bacterial symbionts can help process adult diet components^15^, whereas symbionts in larvae and food stores protect against pathogenic fungi^16^. While highly eusocial bees are rare, stingless bees (*Meliponini*), a sister group to honey bees in the monophyletic Apidae clade^17^, independently evolved complex eusociality, resulting in distinct colony structures and developmental strategies. Unlike *A. mellifera* — which has vertical combs with pollen, honey and brood (fed daily) — many stingless bees have horizontal brood combs separate from pollen and honey pots, and larvae receive their entire diet in a single provisioning before the egg is laid (Fig. 1a)^18^. Additionally, many stingless bee species, such as from *Scaptotrigona* genus, depend on a diet complemented with yeast symbiont for successful development^19,20^, making them an ideal system for studying multi-kingdom social microbiome assembly.

**Fig. 1:**
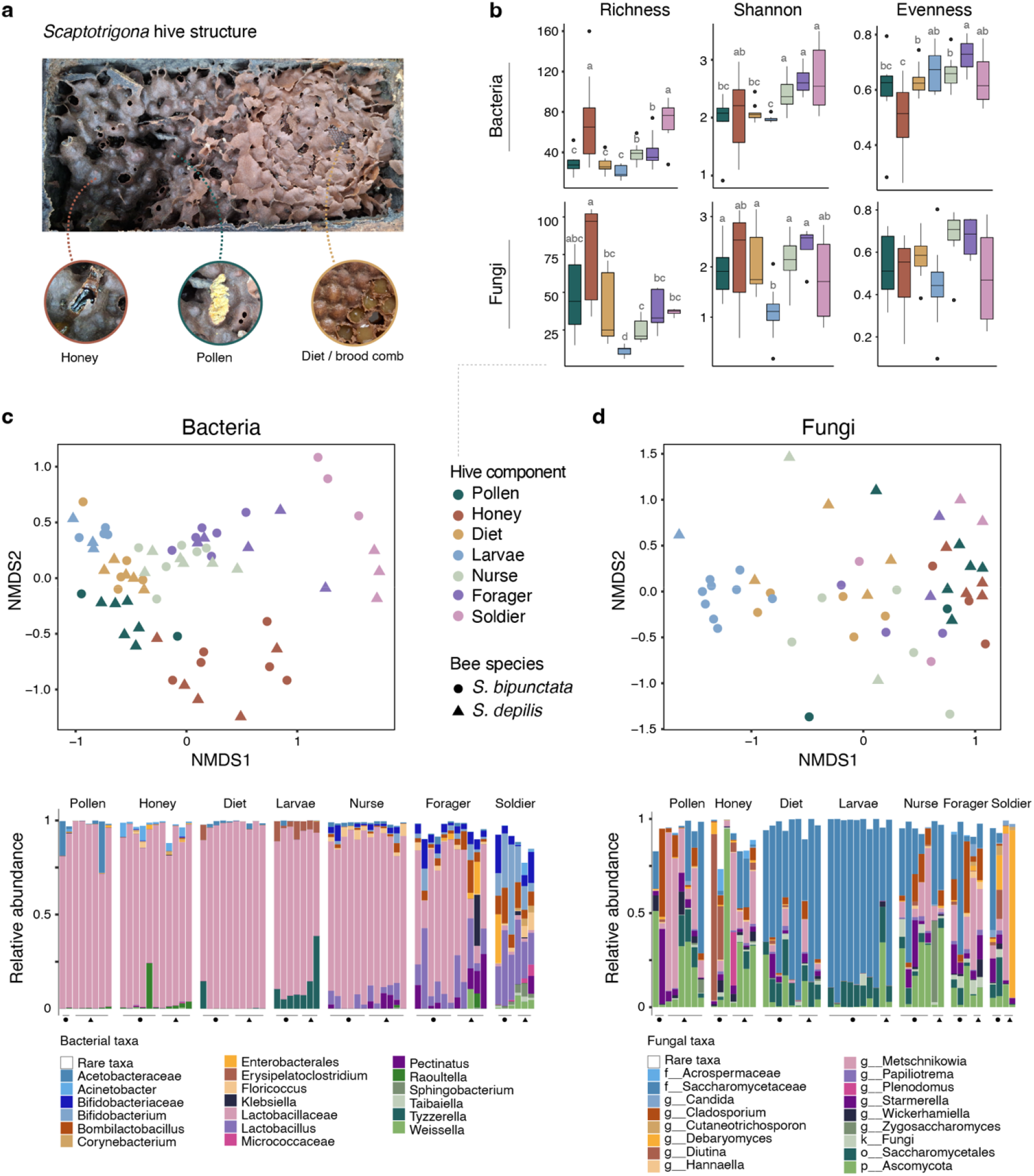
Multi-kingdom microbiome of *Scaptotrigona* stingless bee hive. (**A**) Image with the above view of *Scaptotrigona* hive structure highlighting key differences from other eusocial bee hives. As many stingless bees, *Scaptotrigona* hives have separate pots for pollen and honey storage on one side, while horizontal combs with developing brood on the other side, covered with an additional protective cerumen layer. As shown in the “Diet/brood comb”, each individual brood cell contains the complete diet for larval development, deposited by specialized nurse bees before the queen lays the egg, and is immediately capped afterward. Image credit: Irmgard I. W. Caesar. (**B**) Alpha-diversity metrics of amplicon sequencing variants (ASVs) for bacterial (top) and fungal (bottom) communities across sample types, including pollen, honey, larval diet, larvae, nurse, forager, and soldier bees. Microbiome richness varied significantly (Kruskal-Wallis, Bacteria: χ² = 31.86, df = 6, p = 1.735 × 10 5; Fungi: χ² = 30.58, df = 6, p = 3.05 × 10 5), as did diversity (ANOVA, Bacteria: F = 4.95, p = 5.8 × 10 4; Fungi: F = 2.71, p = 0.033). Differences in evenness were only observed in bacterial communities across sample types (Kruskal-Wallis: χ² = 23.83, df = 6, p = 5.62 × 10 4). Letters above the bars represent differences among samples based on post-hoc test with adjusted p-values, where significance is considered for p < 0.05. (**C**) Beta-diversity analysis shows bacterial community compartmentalization across hive components (PERMANOVA; Bray-Curtis dissimilarity: pseudo-F = 7.45, df = 6, p = 0.001), with ASV-level differences between *Scaptotrigona* species, despite overall dominance by Lactobacillaceae (bottom barplot). (**D**) The fungal community was also different across samples types (PERMANOVA; Bray-Curtis dissimilarity: pseudo-F = 2.32, df = 6, p = 0.001), specially in larval diet and larvae samples which are primarily colonized by Saccharomycetes, also showing ASV-level differences between *Scaptotrigona* species.

Here, we investigate ecological drivers of microbiome assembly in the highly eusocial stingless bees (*Scaptotrigona depilis* and *S. bipunctata*), an emergent and key system to study microbiome ecology and evolution. Using amplicon sequencing and shotgun metagenomics, we characterize bacterial and fungal communities across hive components, such as food stores, developing larvae, and adult castes — including a distinct subcaste known as the soldier^21^. Our results reveal a multi-layered mechanism of microbiome assembly, spanning host biology to cross-kingdom interactions. We show that the *Scaptotrigona* hive structure and caste/subcaste system contributes to microbiome segregation, with bacteria carrying genes potentially involved in local establishment. Additionally, while fungal communities appear transient in most hive components, a distinct community is maintained in the spatially protected larval diet through cross-kingdom interactions.

## Results

### *Scaptotrigona* hive microbiome is dominated by Lactobacillaceae and Saccharomycetaceae

Current knowledge of *Scaptotrigona* microbiome is limited to culture-dependent studies or amplicon sequencing of forager bees^19,22,23^. To characterize its hive microbiome and determine if a consistent microbial community is associated with the bee genus or species, we performed 16S rRNA (bacteria) and ITS (fungi) amplicon sequencing on two species, *S. depilis* and *S. bipunctata*. Hives from three meliponaries were sampled for various components (**Fig. 1A**), including food stores (pollen and honey), developing bees (larvae and larval diet), and adult bees (worker subcastes: nurse, forager, and soldiers). A total of 691 bacterial amplicon sequencing variants (ASVs) were recovered from 65 samples, and 910 fungal ASVs from 50 samples (**Supplementary Table 1**). ASV richness and diversity varies significantly among hive components (**Fig. 1B**). Both bacterial (Kruskal-Wallis: χ^²^ = 31.86, df = 6, p = 1.735 × 10^□5^) and fungal (Kruskal-Wallis: χ^²^ = 30.58, df = 6, p = 3.05 × 10^□5^) microbiomes vary in richness.

Honey and soldier bees, similar in age to foragers but tasked with nest defense^21^, have the richest microbial communities, especially for bacteria. Diversity also differs significantly when comparing the fungal microbiome between hive components (ANOVA: F = 2.71, p = 0.033), but particularly for the bacterial community (ANOVA: F = 4.95, p = 5.8 × 10^□4^), with adult bees having the highest diversity. Bacterial evenness is higher in adult bees, larvae, and larval diet (Kruskal-Wallis: χ^²^ = 23.83, df = 6, p = 5.62 × 10^□4^), despite lower richness and diversity in the latter.

When comparing bacterial microbiome composition (**Fig. 1C**), hive component is the most influential factor (PERMANOVA; Bray-Curtis dissimilarity: pseudo-F = 7.45, df = 6, p = 0.001), as shown by the distribution of samples in the non-metric multidimensional scaling analysis. Most hive components are dominated by Lactobacillaceae, except for soldier bees, with less abundant taxa contributing to community differences. For example, food stores also carry Acetobacteriaceae and *Acinetobacter*, while adult bees host *Floricoccus*, *Bifidobacteriaceae*, and *Bombilactobacillus*. Differences in microbiomes of some hive components, such as soldier bees and honey, are also explained by the meliponary (PERMANOVA; Bray-Curtis dissimilarity: pseudo-F = 1.34, df = 11, p = 0.026), possibly due to constant environmental exposure. The fungal community (**Fig. 1D**) also shows significant differences among hive components (PERMANOVA; Bray-Curtis dissimilarity: pseudo-F = 2.32, df = 6, p = 0.001). Unlike the bacterial community, these differences are mostly explained by larvae and larval diet. While food stores and adult bees harbor a more variable fungal composition, larval diet and larvae microbiomes are predominantly comprised of Saccharomycetaceae. This group may include unclassified *Zygosaccharomyces* and *Starmerella* — all known associates of *Scaptotrigona* bees^24^ and potentially underrepresented in public ITS databases. There are no differences for bacterial or fungal composition between bee species by hive components (two-way PERMANOVA; Bray-Curtis dissimilarity, Bacteria: pseudo-F = 1.0730, df = 6, p = 0.4; Fungi: pseudo-F = 1.1837, df = 6, p = 0.09).

### Segregation of bacterial species and limited strain sharing across hive components

To gain deeper insights into the microbial species and strains in a *Scaptotrigona* hive, we performed shotgun sequencing on sample pools from two hives of each stingless bee species within a single meliponary. The 28 metagenomes yielded 9 to 28 million paired reads. By mapping reads to SSU barcode regions we see that the *Scaptotrigona* microbiome is primarily bacterial, with fungi present in much lower proportions (**Supplementary** Fig. 1) — thus no metagenome-assembled genomes (MAGs) of fungi were recovered. We recovered 14 high-quality bacterial MAGs (**Supplementary Table 2**) after dereplicating at species level; representing prevalent bacteria identified in our amplicon data. Nine MAGs are from bacteria consistently associated with at least one hive component (coverage >1X, breadth >50%): three *Apilactobacillus*, two *Bombilactobacillus*, one *Acinetobacter*, *Bifidobacterium*, *Raoultella*, and *Wolbachia pipientis* (**Fig. 2A**). Except for *Wolbachia*, these MAGs represent novel species with average nucleotide identity (ANI) ranging from 67% to 75% to their closest relatives. The *Wolbachia* MAG had 96% ANI with its closest relative, *Wolbachia* wLug. Combined with its detection only in soldiers (**Fig. 2B, Supplementary Table 1**), this suggests that *Wolbachia* DNA may reflect contamination from recent interactions with hive parasites carrying the strain.

**Fig. 2:**
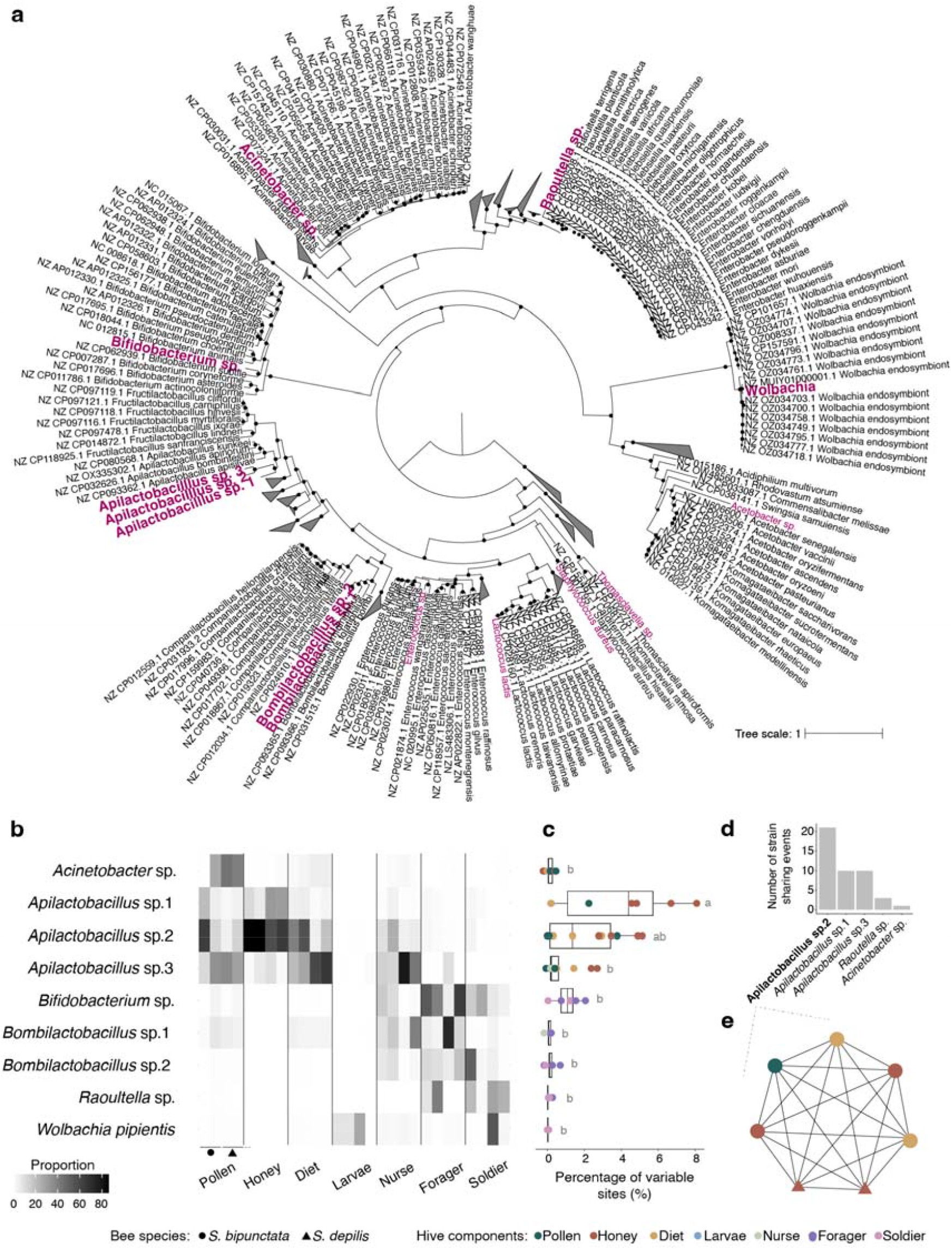
Bacterial species and strains in *Scaptotrigona* hives. (**A**) Maximum likelihood phylogenetic analysis of high-quality metagenome-assembled genomes (MAGs), based on the alignment of 73 single-copy orthologs present in >80% of 500 reference genomes. MAGs from *Scaptotrigona* hives are highlighted in pink, with the nine species consistently associated with hives (appearing in >1 sample, >1× coverage, and >50% breadth) shown in bold and included in subsequent analyses. Dots indicate nodes with >95% bootstrap support. (**B**) Proportions of bacterial species in hive components for each species, shown in order: two *S. bipunctata* and two *S. depilis* samples. To avoid misleading interpretations, for larval microbiomes — which are only minimally colonized — coverage proportions refer to the proportion to all MAGs mapped (see Fig. S2). (**C**) Differences in the percentage of variable sites per bacterial species (Kruskal-Wallis χ² = 37.51, df = 5, p = 4.73 × 10 7), shown as boxplots with points representing individual samples. Letters above bars represent significant differences based on post-hoc tests with adjusted p-values (p < 0.05). Hive components (colors) also show differences in strain diversity (Kruskal-Wallis χ² = 20.75, df = 5, p = 0.0009). (**D**) Bacterial species with strains shared at least once between sample pairs (e.g., sample A and sample B carry the same strain), with bars indicating the number of sharing events. (**e**) *Apilactobacillus* sp.2 strain-sharing network, highlighting hive components and species carrying the same strain.

Among MAGs in the same genera, the two *Bombilactobacillus* share 87% ANI, while the three *Apilactobacillus* share 67–78% ANI. Their closest related strains, *B. thymidiniphilus* and *A. apisalvae*, are from stingless bees native to Australia^25^.

Bacterial species exhibit clear segregation within the hive (**Fig. 2B**), also corroborating findings from amplicon sequencing. Food stores and larval diet are primarily inhabited by Lactobacillaceae, predominantly the phylogenetically derived strains *Apilactobacillus* sp. 2 and *Apilactobacillus* sp. 3, as well as Acetobacteraceae and *Acinetobacter sp.* Interestingly, *Apilactobacillus* sp. 2 dominates the *S. bipuctata* larval diet, while *Apilactobacillus* sp. 3 dominates in *S. depilis*. Larvae are minimally colonized by bacteria, with no MAG coverage above 1X depth or 50% breadth, and non-persistent hive strains likely reflecting transient environmental contamination (**Supplementary** Fig. 2). Following development, the adult bee microbiome undergoes a transition. Nurse bees, young workers tasked with feeding and caring for the brood, host *Apilactobacillus* — especially strains abundant in larval diet — along with *Bombilactobacillus* (Lactobacillaceae) and *Bifidobacterium* (Bifidobacteriaceae). Forager bees, older workers responsible for collecting nectar, pollen, water, and propolis, no longer carry *Apilactobacillus* strains but maintain *Bombilactobacillus* and *Bifidobacterium*. Nest entrance- defending soldier bees are mostly colonized by *Bifidobacterium*. However, some also carry Enterobacteriaceae, such as *Raoultella* sp., related to opportunistic pathogens^26^ and, as *Wolbachia*, may reflect transient environmental contamination acquired through interactions with intruders.

Among hive components, honey, pollen, and forager bees serve as the primary entry points for bacterial strains, showing the highest strain diversity, as indicated by the percentage of variable sites in MAGs (**Fig. 2C**; Kruskal-Wallis χ² = 20.75, df = 5, p = 0.0009). *Apilactobacillus* sp. 3, the least abundant of the three *Apilactobacillus* species, has the greatest strain diversity, followed by *Apilactobacillus* sp. 2, both predominantly associated with food stores (Kruskal-Wallis χ² = 37.51, df = 5, p = 4.73 × 10^□7^). The larval diet, a mixture of pollen and nectar/honey prepared by nurse bees, shows lower strain diversity compared to honey and some pollen samples. Despite overlapping ASVs and bacterial species, strain sharing is rare, with no *Bombilactobacillus* or *Bifidobacterium* strains shared among adult bees — distinct strains dominate each individual. Strain-sharing is mostly seen in food store samples (**Fig. 2D, E**), where, for example, the same *Apilactobacillus* sp. 2 strain is detected across pollen, honey, and larval diet. This strain is also present in both *Scaptotrigona* species, suggesting priority effects may contribute to the dominance of bacterial lineages in each host.

### Bacteria carry genes to handle toxins and degrade plant polysaccharides

To further understand microbiome assembly dynamics in *Scaptotrigona* hives, we functionally annotated all bacterial contigs from each sample, alongside the eight MAGs consistently associated and representing novel bacterial species. The bacterial contig annotations confirm that larvae are minimally colonized by bacteria, with the total length of bacterial contigs in larvae samples ranging from 0.01 to 0.05 Mb, while adult bees range from 2 to 13 Mb and food stores from 14 to 33 Mb. Fewer than three proteins from larvae samples were classified into functional groups, leading us to exclude those samples from further analysis (**Supplementary Table 3**). Hive components differ significantly in the number of predicted proteins from bacterial contigs (ANOVA: F = 3.04, p = 0.037). The predicted proteins were classified into KEGG Orthology (KO) groups and organized into BRITE functional hierarchies. On average, 45% (± 6%) of proteins were assigned to KOs, with over 99.5% of these KOs linked to BRITE categories. The overall number of bacterial proteins positively correlates with expanded functional capacity, indicated by a greater number of unique KOs (Pearson correlation: r = 0.56, p = 0.01; **Fig. 3A**). However, this correlation is not significant when we include diet samples (Pearson correlation: r = 0.33, p = 0.12), which have high protein counts but low KO classification (around 40%; **Supplementary Table 3**) — suggesting the presence of bacteria carrying novel functions in this environment.

**Fig. 3:**
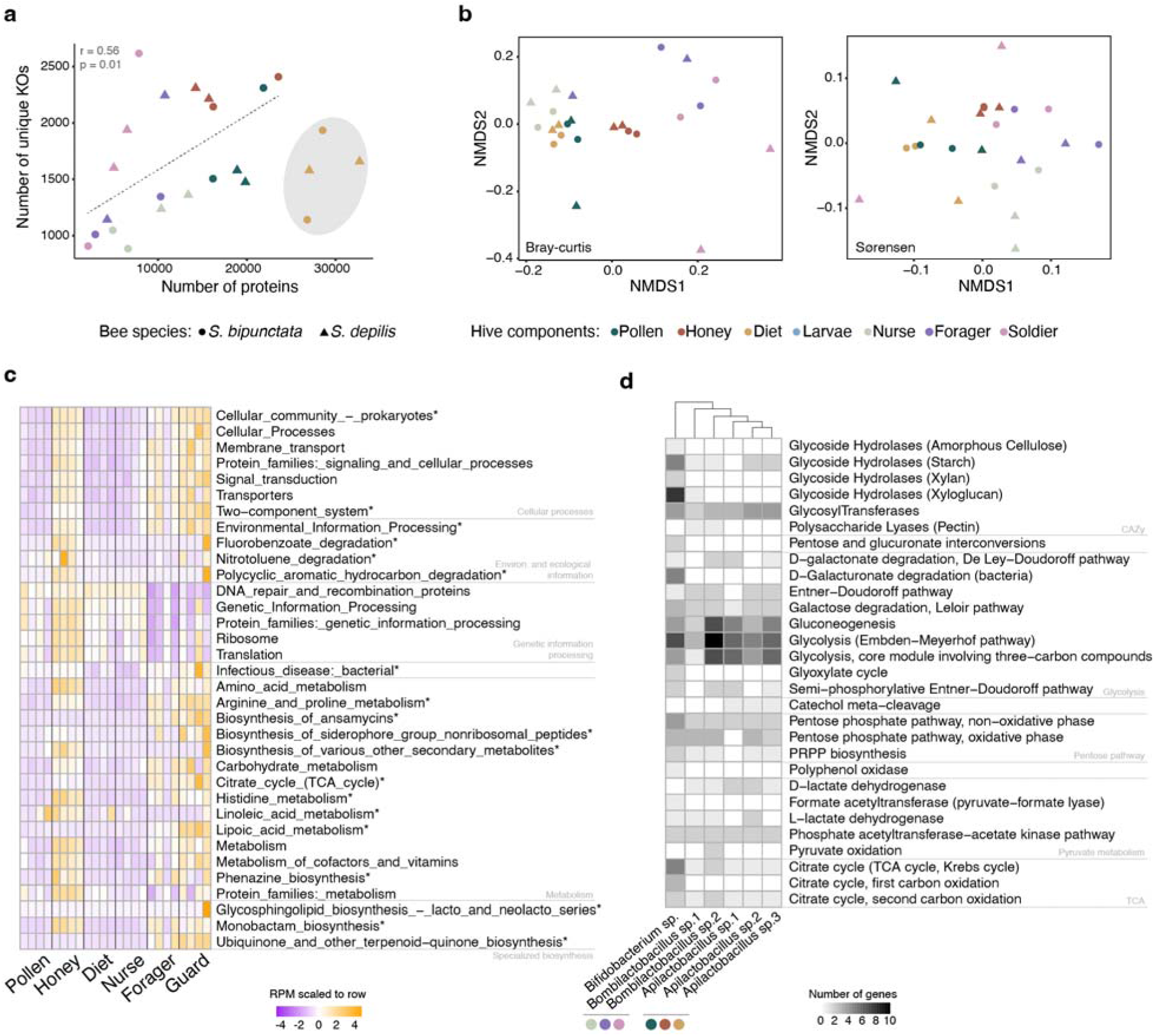
Functional repertoire of bacteria across *Scaptotrigona* hive components. (**A**) Number of proteins and non-redundant KEGG Orthology (KO) terms predicted from bacterial contigs in each sample per hive component (color) and species (shape). A positive Pearson correlation between protein count and number of unique KOs was observed only when larval diet samples (highlighted in grey) were excluded due to their lower KO assignment rates (Table S3). (**B**) Non-metric multidimensional scaling (NMDS) analysis of samples by species (shape) and type (color) based on normalized coverage of BRITE categories using Bray-Curtis dissimilarity and presence/absence using Sørensen distances. (**C**) Normalized coverage of BRITE categories per sample, displaying the top 15 most abundant categories across all samples (Table S4) and the top 10 categories enriched in guard and honey samples (* = adjusted p < 0.05; Table S5). Coverage values are row-scaled for better visualization. (**D**) Number of genes involved in different carbon metabolism pathways across the six most prevalent bacterial species in *Scaptotrigona* hives, with columns clustered based on Euclidean distance. Colored circles at the bottom indicate the hive components where these species are most prevalent – following color key in (**A, B**).

Comparisons of functional categories show that larval diet, nurse bees, and pollen have some overlap, while honey, forager, and soldier samples exhibit more distinct functional profiles (**Fig. 3B, Supplementary Table 4**). Indeed, honey and soldier samples have more unique KOs and are the only hive components with significantly enriched BRITE categories (**Fig. 3C, Supplementary Table 5**). Soldiers, for example, show enrichment in bacterial infectious disease proteins and ansamycin biosynthesis (Wilcoxon rank-sum test, adjusted-p = 0.04, fold-change = 1.75; adjusted-p = 0.04, fold-change = 7.77), which can be coming from pathogens or being used against them. Honey is enriched in proteins related to antimicrobial compound biosynthesis, such as phenazines and monobactams (Wilcoxon rank-sum test, adjusted-p = 0.03, fold-change = 4.22; adjusted-p = 0.01, fold-change = 4.29), with potential roles in competition and spoilage prevention. Honey is also enriched for proteins involved in plant-derived compounds detoxification like nitrotoluene, fluorobenzoate, and polycyclic aromatic hydrocarbon degradation (Wilcoxon rank-sum test, adjusted-p = 0.02, fold-change = 6.51; adjusted-p = 0.01, fold-change = 5.35; adjusted-p = 0.01, fold-change = 6.32). The hive microbiome overall shows relative high abundance of genes for carbohydrate and amino acid metabolism, as well as cofactors and vitamins metabolism (**Fig. 3C**). For bacterial species prevalent in each hive component, we additionally predicted carbon metabolism (**Supplementary Table 6**).

*Apilactobacillus*, a lactic acid bacterium (e.g., D-lactate dehydrogenase), is abundant in food stores and may contribute for spoilage control through fermentation. In contrast, *Bifidobacterium* and *Bombilactobacillus*, which are abundant in adult bees, carry more genes for plant polysaccharide digestion (e.g., glycoside hydrolases, polysaccharide lyases targeting pectin), potentially contributing to bee nutrition (**Fig. 3D**).

### Cross-kingdom interactions shape the characteristic brood microbiome

The functional analyses revealed bacterial adaptations to hive components, including potential roles in adult bee metabolism. In non-bee hive components, bacteria likely persist due to their ability to thrive in high-sugar and fermented environments (**Fig. 3D**). However, we described a consistent bacterial and fungal community especially in larval diet — where microbial interactions may play a significant role on assembly once there is no bee innate immune system or extra host selective pressures. We thus wondered how bacteria interact with fungi in this hive component, especially since *Scaptotrigona* depends on the yeast *Zygosaccharomyces* for its development^27^. The dominant bacteria in the larval diet, *Apilactobacillus*, is closely related to taxa with known antifungal role^28,29^ — suggesting that bacteria in stingless bee larval diet may not inhibit fungi or may exhibit selective interactions. To investigate such interactions, we isolated microbes from larval diet **(Supplementary Table 7**), and conducted *in vivo* and *in vitro* experiments. Three bacterial strains were tested: *Apilactobacillus*, the dominant bacteria in the larval diet; *Weissella* (Lactobacillaceae), rare in the larval diet but found in food stores and older workers (**Fig. 1C**); and *Bombella* (Acetobacteriaceae), also rare (**Supplementary Table 1**) but from the same genus as a key honey bee larval symbiont with antifungal role^16^. Our competition assay shows that all strains inhibit the bee pathogen *Aspergillus* (**Fig. 4B**) — confirmed to be pathogenic in our honey bee larval assays (**Supplementary** Fig. 3). *Apilactobacillus* exhibits the strongest inhibition, preventing both sporulation and filamentous growth. This left us with a conundrum: if these bacteria share larval diet with the symbiotic yeast, do they also inhibit their growth *in vitro*?

**Fig. 4:**
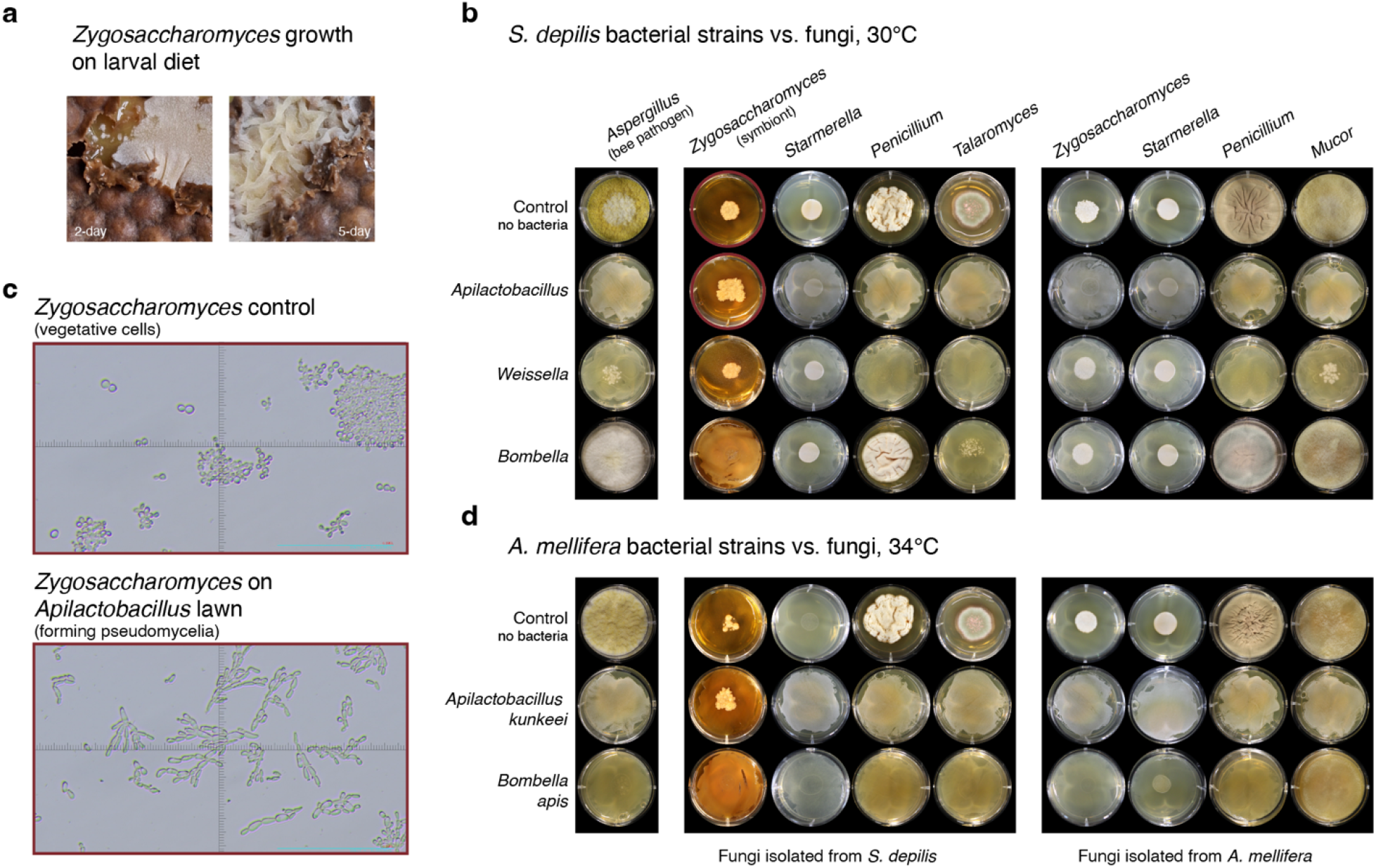
Competition assays between bee development-associated bacteria and fungi. (**A**) Growth of *Zygosaccharomyces* in brood combs brought to the laboratory after egg removal to allow yeast proliferation, which larvae would typically consume post-hatching. Images taken at 2 and 5 days. Image credit: Irmgard I. W. Caesar. (**B**) Competition assays between bacterial isolates from *Scaptotrigona depilis* larval diet and fungal isolates from the same environment, as well as fungal isolates from *Apis mellifera* larvae. The experiment included the stingless bee symbiont *Zygosaccharomyces* and the bee pathogenic fungus *Aspergillus*. The control row shows fungal growth (15 μL MRS media + 1000 spores or 5 μL yeast at OD600 1) without a bacterial lawn. Subsequent rows show fungal growth on a bacterial lawn (15 μL of 10 CFU). (**C**) Light microscopy images of 15-day Zygosaccharomyces cells grown without bacteria (control; above) and on an *Apilactobacillus* lawn (below). Cyan scale bar = 0.25 μm. (**D**) Competition assays between bacterial isolates from *A. mellifera* larvae and fungal isolates from both *S. depilis* larval diet and *A. mellifera* larvae, using the same plating design as described in (**B**).

We next tested whether these bacteria from *S. depilis* larval diet could inhibit fungi isolated from the same environment, including the symbiont *Zygosaccharomyces, Starmerella* yeast, and the filamentous fungi *Talaromyces and Penicillium* (**Supplementary Table 7**) — only the latter potentially pathogenic based on honey bee larval assays (**Supplementary** Fig. 3). Our results show that while the two Lactobacillaceae mostly inhibit the filamentous fungi to varying degrees, *Bombella* inhibits *Zygosaccharomyces* (**Fig. 4B**). Interestingly, *Apilactobacillus* does not inhibit *Zygosaccharomyces* and promotes its growth, accelerating pseudomycelium formation compared to controls without a bacterial lawn (**Fig. 4B, C**) — structure suggested to support its floating in larval diet (**Fig. 4A**)^27^. *Apilactobacillus* also promotes *Zygosaccharomyces* growth even in the presence of *Bombella* (**Supplementary** Fig. 4). We also tested for bacterial-load effects with experiments using lower bacterial CFUs and observed similar results (**Supplementary** Fig. 5).

### *Apilactobacillus* inhibits honey bee associated fungi

To assess the spectrum of such selective antifungal role, particularly from *Apilactobacillus*, we tested how these bacteria would interact with fungi not typically found in stingless bee hives. Using an eco-evolutionary framework, we isolated fungi from *Apis mellifera* larvae, which share a recent ancestor with stingless bees^17^ and may exchange microbes via shared floral resources^30^. The tested fungi included *Zygosaccharomyces*, *Starmerella*, and the filamentous fungi *Penicillium* and *Mucor* (**Supplementary Table 7**). Previous studies suggest all these fungi are likely transient, and our *in vivo* assays indicate that *Penicillium* is potentially pathogenic to the larvae (**Supplementary** Fig. 3). *Apilactobacillus* and *Weissella* from *Scaptotrigona* inhibit the filamentous fungi but not *Starmerella*. Interestingly, while *Apilactobacillus* promotes *Zygosaccharomyces* from *S. depilis*, it inhibits the honey bee-associated *Zygosaccharomyces* (**Fig. 4B**), suggesting a highly selective positive interaction between the stingless bee-associated microbes.

Finally, we conducted inhibition assays with laboratory strains of *A. kunkeei* and *B. apis*, predominant in honey bee larvae (**Supplementary Table 7**). Both bacteria inhibit the growth of all filamentous fungi (**Fig. 4D**), confirming their defensive role in honey bees. Interestingly, *A. kunkeei* exhibits a similar selective inhibition pattern to *Apilactobacillus* from *S. depilis*, promoting only the growth of stingless bee-associated *Zygosaccharomyces*. This suggests that *Zygosaccharomyces* in *Scaptotrigona* may also have evolved strategies to counteract *Apilactobacillus*. We considered brood temperature differences between bee species for our experiment, which end up revealing that *S. depilis* yeast also grow optimally at lower temperatures (∼30°C) typical of their hives^31^, compared to ∼34°C for *A. mellifera*^32^ — indicating temperature as an additional selective pressure shaping the fungal microbiome during bee development.

## Discussion

Microbiome assembly and maintenance are critical for hosts that depend on their microbiomes for health and fitness. Eusocial insects, as superorganisms, have direct fitness determined by collective colony performance^14^, adding an extra layer of complexity to the study of their microbiomes. We present the first comprehensive microbiome characterization of multiple components of the highly eusocial insect colony, *Scaptotrigona* stingless bees, using bacterial and fungal amplicon sequencing, metagenomics, and experiments with isolated microbes. We show that *Scaptotrigona* harbors a stable microbiome dominated by Lactobacillaceae and Saccharomycetaceae, drawing an interesting parallel to many other systems with similar taxa. This microbial community is spatially segregated across hive components, with novel species, carrying genes potentially involved in local adaptation. Particularly in the brood component of the hive, we show the larval diet hosts a stable Saccharomycetaceae community, sustained by brood comb features and highly selective interactions with *Apilactobacillus*. These findings highlight the role of multi-layered mechanisms in shaping microbiome assembly and maintenance, from host biology to cross-kingdom interactions (**Fig. 5**).

**Fig. 5:**
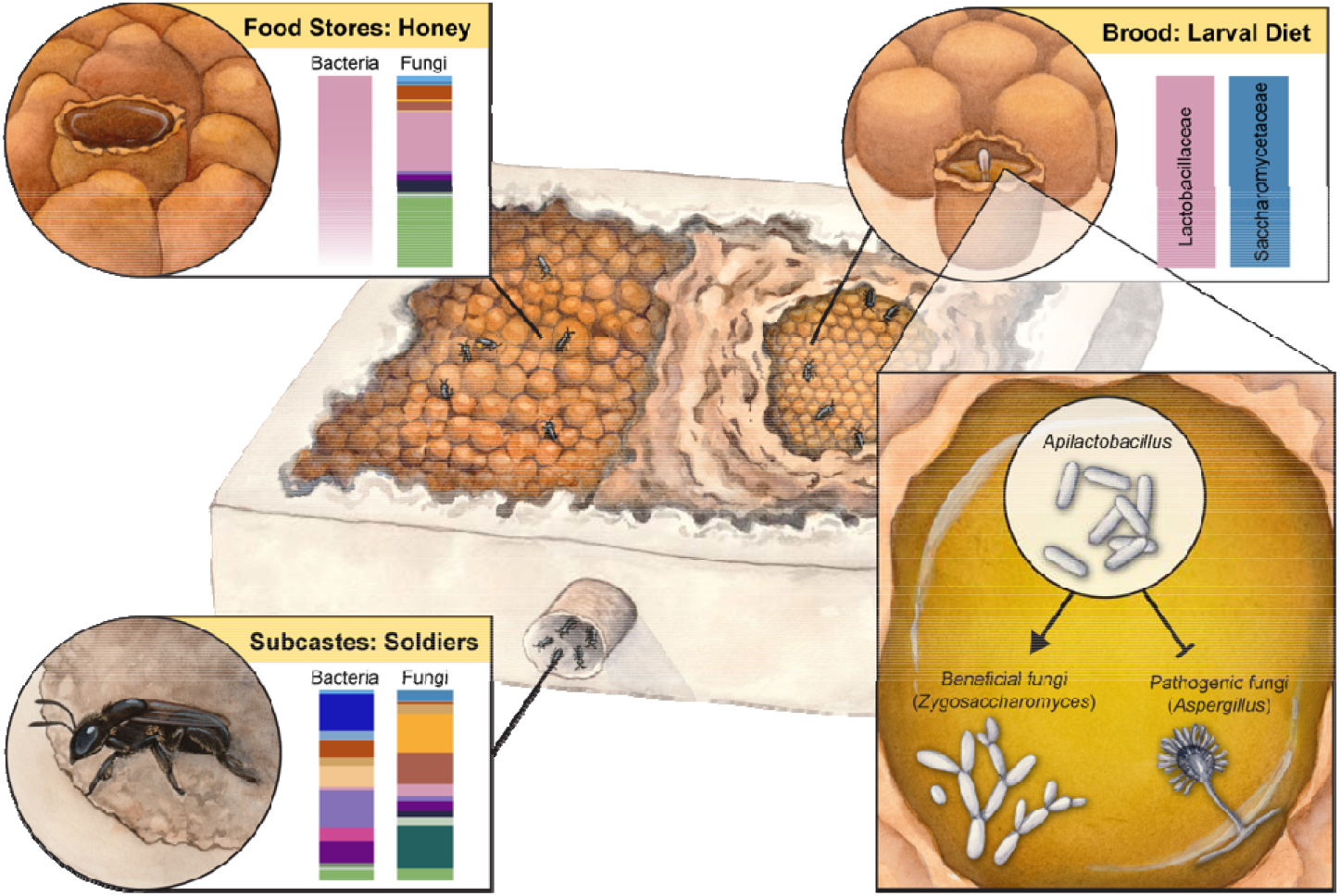
Multi-level mechanisms in stingless bee microbiome assembly. Many mechanisms likely shape colony microbiomes, and based on our results, we discuss at least two key ones in stingless bee hives: host biology and cross-kingdom microbial interactions — both potentially coevolving to promote and protect one of the most important components of a social insect colony, the developing brood and its microbiome. Our findings of microbial segregation within spatially distributed hive components (e.g., food stores and subcastes) suggest greater opportunities for local adaptation as well as better control of microbial influx. Hive components apart from the brood combs show high bacterial strain diversity (see Honey, pink gradient) and carry many transient bacteria and fungi (see Soldiers) potentially pathogenic. Below all the physical layers protecting the brood, our results also reveal a consistent community dominated by Lactobacillaceae and Saccharomycetaceae, which is not only assembled as a consequence of the hive biology but also through selective bacteria-fungi interactions we observed with in vitro assays—with *Apilactobacillus* promotes the growth specifically of the beneficial symbiotic yeast while inhibiting other fungi, such as potential fungal pathogens. Illustration(s) by Julie Johnson (Life Science Studios).

Social interactions can homogenize microbiomes through reinoculation, as seen in group- living spiders^33^ and wild lemurs^34^. In eusocial bees, hive connectedness — driven by worker movement and care behaviors — may create similar patterns. In honey bees, for example, larvae, royal jelly, and food stores share the same comb and are all colonized by *A. kunkeei* and *B. apis*^35^. *Scaptotrigona*, however, has a more compartmentalized hive structure which may be contributing to microbial segregation. We show that pollen and honey harbor more *Apilactobacillus* strains, with addition of *Acetobacteraceae* in pollen, while the larval diet supports a multi-kingdom community with fewer *Apilactobacillus* strains and *Saccharomycetaceae.* Additionally, social organization with subcastes, along with compartmentalization from metamorphosis, may further regulate microbial exchange and turnover. While it has been already suggested that metamorphosis likely facilitates gut microbiota shifts that support ecological niche transitions^36^, we highlight that the additional compartmentalization within the hive may also facilitate microbial adaptation. Our functional analyses show that bacteria in the larval diet promote yeast growth and harbor genes associated with tolerance to high-sugar and acidic conditions, potentially preventing spoilage, while bacteria associated with adult bees may contribute to the digestion of plant-derived diets.

Stingless bee hive structure and biology also seem to limit environmental microbial influx, contributing for a stable microbiome^37,38^. Using amplicon and strain-level analyses we identify honey as the most microbial rich hive component, and its isolation from the brood may help protect the brood microbiome. The repeated evolution of a soldier subcaste in stingless bees, likely to counter hive invaders^21^, also aligns with our finding of higher bacterial richness in this subcaste. Environmental microbes, including the potential pathogen *Raoultella* sp.^26^, were mostly found on soldiers.

Beyond spatial organization, stingless bee brood combs are enclosed by a waxy cover (cerumen involucrum), and brood cells are capped immediately after egg laying over the larval diet — component hosting the most consistent multi-kingdom microbial community. While the presence of *Zygosaccharomyces* and other yeasts was expected^20,22,39,40^, we provide the first culture-independent evidence of their distribution. Other hive components, like in honey bees, lack a stable fungal community, suggesting transient associations^41,42^. These brood protective layers likely represent an important evolutionary adaptation in eusocial bees, as stingless bees uniquely evolved mass provisioning strategy to feed larvae, in contrast to honey bees or bumble bees, which use progressive provisioning^43^. While this provisioning strategy may have contributed to the establishment of beneficial diet microbes, it may also have selected for mechanisms to prevent the proliferation of harmful microbes. Indeed, we show that cross-kingdom interactions also drive larval diet microbiome assembly and maintenance: *Apilactobacillus* selectively promotes the stingless bee yeast symbiont *Zygosaccharomyces* and inhibits potentially pathogenic fungi, including non-native *Zygosacharomyces* from honey bees.

Selective bacterial-fungal interactions have been observed in other systems, such as fungus-growing ants^44^. The mechanism behind the interaction we describe here requires further investigation, but our results suggest *Apilactobacillus* fungal inhibition may be ancestral, since we observed similar patterns for *A. kunkeei* from honey bees. Nevertheless, while *S. depilis* and *S. bipunctata* share similar microbiomes, our discovery of novel bacterial species and yeasts clustering with stingless bee-derived strains warrants further investigation into Meliponini microbiome codiversification. It has been suggested that the swarming behavior of stingless bees — unique among social insects —, where workers bring materials from the mother nest to the new one, may promote the vertical transmission of *Zygosaccharomyces* and its codiversification with stingless bee species^27^. The same may apply to microbes in other hive components, as well as to prevalent microbes in the larval diet, such as *Starmerella* and *Apilactobacillus*. Indeed, we show that *Apilactobacillus* only weakly inhibits *Starmerella*, a yeast known to promote *Zygosaccharomyces* growth^45^ and which we show to be more similar to another stingless bee strain — due to the limited availability of sequences, we are still unable to observe a clear phylogenetic clustering pattern for *Starmerella* as seen for *Zygosaccharomyces* **(Supplementary** Fig. 6). Additionally, *Scaptotrigona* yeasts thrive at the lower temperatures, characteristic of *Scaptotrigona* hives. Altogether, these findings suggest a tight coevolutionary dynamic between brood combs, larvae, larval diet, bacteria and fungi, warranting further investigation. They also highlight the importance of considering the entire colony when investigating codiversification patterns in eusocial insects.

Our study provides new insights into microbiomes assembly and maintenance, emphasizing the roles of host biology, spatial organization, and cross-kingdom microbial interactions within a eusocial insect colony. The stingless bee microbiome serves as a valuable model for studying host-microbiome ecology and evolution, with broader implications for systems that share similar microbial communities or biological parallels, such as plant tissues and human social networks. Finally, our findings also contribute to a deeper understanding of pollinator health, which is increasingly threatened by agrochemical use and climate change.

## Methods

### Sampling from managed stingless bee hives

Samples of *S. depilis* and *S. bipunctata* were collected from three locations in Rio Grande do Sul, the southernmost state of Brazil: BP (29.4926° S, 51.3550° W), BG (29.1667° S, 51.5170° W), and SC (29.5911° S, 51.3761° W). Sampling was carried out in April 2021, under SISGEN collection permission no. AE1AFA6. The hives were maintained in meliponaries alongside other stingless bee species, without interventions, such as the use of antibiotics, diet supplementation, or heaters. Two hives per species were sampled for honey (from 3 pots), pollen (from 3 pots), larval diet (from 1 egg comb), 3^rd^ instar larvae, nurses, foragers and soldiers. All samples were transported to the laboratory on ice and processed on the same day for microbial isolation or stored at -80°C for subsequent DNA extraction.

### Amplicon and metagenome sequencing

DNA was extracted from each sample using the PureLink™ Genomic DNA Mini Kit (Thermo Fisher, K182001), following the manufacturer’s protocol with an additional initial lysis step using Lysozyme (Thermo Fisher Scientific, 89833). Honey (3 mL) and larval diet (300 µL) samples were first diluted with 1 volume of 1X PBS, vortexed, then centrifuged at 8,000 rpm for 10 min, and the pellet used for DNA extraction. For adult bees, the guts of 7 individuals were dissected using sterile forceps and pooled. Similarly, 7 entire 3^rd^ instar larvae were pooled for DNA extraction. DNA quality was assessed using the Genomic DNA (gDNA) ScreenTape assay on an Agilent 2200 TapeStation (DIN > 7 cutoff). DNA concentration was measured with the Qubit dsDNA HS kit (Invitrogen) on a ThermoFisher Qubit4 fluorometer. To control for possible contaminants, we used blank extractions and sterile water as negative controls during the PCR.

PCR amplification was performed using 16S rRNA and ITS primers from the Earth Microbiome Project^46^, with the Illumina adapter, a primer pad, and a linker added to the 5’ end, and a barcode added to one primer direction. The V4 region of the 16S SSU rRNA was targeted using primers 515F (GTGYCAGCMGCCGCGGTAA) and 806R (GGACTACNVGGGTWTCTAAT), generating an amplicon of ∼390 bp ^47,48^. For the ITS region, primers ITS1 (CTTGGTCATTTAGAGGAAGTAA) and ITS2 (GCTGCGTTCTTCATCGATGC) were used, generating an amplicon of ∼260 bp ^49,50^. PCR reactions were performed in triplicate, with each 25 µL reaction containing 2 µL of template DNA (100 ng total), 12.5 µL of 2X Phusion® High-Fidelity PCR Master Mix with HF Buffer (NEB, M0531L), and 0.75 µL of each primer at 10 µM. For 16S rRNA, the PCR conditions included an initial denaturation at 98°C for 3 min, followed by 30 cycles of denaturation at 98°C for 20 s, annealing at 71°C for 30 s, and extension at 72°C for 45 s, with a final extension at 72°C for 10 min. For ITS, the annealing was performed at 54°C for 35 s, and the extension at 72°C for 7 s. The triplicates were pooled, and the amplicons were first checked on a 1.5% agarose gel, then quantified using the Quant-iT™ PicoGreen™ dsDNA Assay Kit (Invitrogen, P7589) before pooling. The 16S and ITS libraries were prepared using the TrueSeq Nano Library Preparation Kit (Illumina) and paired-end sequenced (2 × 250 nt) on an Illumina MiSeq flowcell with the MiSeq Nano Kit v2 (500 cycles).

The DNA samples from BP meliponary were also used for shotgun metagenome sequencing. Libraries were prepared using the Illumina DNA Prep kit and IDT 10 bp UDI indices and sequenced on an Illumina NextSeq 2000, producing 2 × 151 bp reads.

Demultiplexing, quality control, and adapter trimming were performed with bcl-convert (v3.9.3) from Illumina. Raw shotgun sequencing data can be found within the NCBI BioSample accessions: SAMN46439779, SAMN46439780.

### Amplicon data and statistical analyses

Illumina fastq files were demultiplexed, barcodes removed and concatenated into a separated barcode files directly after sequencing. The following analyses were conducted with DADA2. As part of the read quality control step, 16S rRNA reads were filtered other than the default using truncLen=c(200,225) and minLen=175, and ITS reads with truncLen=c(190,140), minLen=100. After, reads were dereplicated, inferred ASVs, merged, and chimeric ASVs were removed. The database SILVA v.138.1 (silva_nr99_v138.1_train_set.fa and silva_species_assignment_v138.1.fa) and UNITE v.9 (sh_general_release_dynamic_25.07.2023) were used for the taxonomic assignment of the ASVs. Contaminants in the samples found in the negative controls were removed using decontam R package^51^. We additionally removed any ASVs identified as mitochondria or chloroplasts. The ASVs and related data can be found in **Supplementary Table 1**.

All statistical analyses and figure plots were performed in R version 4.3.1. To describe alpha diversity in the samples we use the functions *estimate*, *diversity* and *specnumber* from the package Vegan^52^. To test for differences between samples and groups, the estimates with normal distribution according *shapiro.test* were compared using t test or ANOVA. In case of no normal distribution, we used Wilcoxon rank sum test or Kruskal-Wallis rank sum test. Following statistical significant Wilcoxon rank sum test we used *pairwise.wilcox.test* as posthoc test. To investigate beta diversity, we first calculated Bray-Curtis and Sørensen dissimilarity dissimilarities and a non-metric multidimensional scaling (NMDS) using *vegdist* and *metaMDS* function from Vegan^52^. We also used these dissimilarity matrices to carry out PERMANOVA (999 permutations) with function *adonis2*. All the results were visualized and plotted with ggplot2 package^53^.

### Metagenome assembly and binning

Paired-end raw sequencing reads from the 28 samples from BP were first trimmed by length and quality using Trimmomatic v.0.36^54^, with options *LEADING:28 TRAILING:28 SLIDINGWINDOW:6:25 MINLEN:75*, then assembled into contigs with metaSPAdes v.3.15.5^55^ using default settings. Only contigs >1500 bp were used for the following steps. Contig coverage was recovered by mapping trimmed individual sample reads back to all contigs with Bowtie2 v.2.4.2^56^, using the options *--no-discordant --no-mixed*. Next, we used Metabat v.2.15^57^ for binning, with flag *-m 1500*, followed by CheckM v.1.2.2^58^ to recover bin quality. Bins with >80% completeness and <5% contamination were dereplicated with dRep^59^ *dereplicate, --P_ani 0.95 --S_ani 0.95 --cov_thresh 0.85*. No fungal bins were recovered, with 99% of bins having <5% completeness according to BUSCO, using fungi_odb10 (2024-01-08, number of genomes: 549, number of BUSCOs: 758) and none of those being confirmed as fungi according to best blast hits. Moving on with the bacterial bins, to recover their coverage per sample we used again Bowtie2 v.2.4.2^56^ to map trimmed individual sample reads back to the bins in a single database. Samtools v.1.17^60^ was used for file transformation, the depth command with option *-a* and *idxstats* command were used to recover the number of reads covering each base and the entire contig. Bin average coverage was calculated as the *number of mapped reads*150/bin length*, and proportions (%) plotted. To estimate the approximate ratio of bacteria to fungi, trimmed reads per sample were mapped with Bowtie2 v.2.4.2^56^ reads against SILVA SSU database v.138.1^61^ which contains 2,224,740 SSU rRNA sequences. Samtools v.1.17^60^ was used for file transformation, and idxstats command to recover the number of reads per sequence contig of different taxa. Total number of reads mapping to bacteria, fungi and insecta were plotted.

### Strain diversity metagenome-assembled genomes

To estimate strain richness of MAGs in each sample we used the command *profile* from inStrain v1.4.0^62^, with *--database_mode* flag, on the bam mapping files generated previously (See section “Metagenome assembly and binning”). As part of inStrain protocol, only MAGs with >1x and 50% breach were considered, and only sites covered at least 5x and the allele frequency of at least 0.05 were counted as SNVs. The output with genome information was used to calculate the percentage ratio of the number of single nucleotide variants (SNVs) to the total length of the MAG covered (*percentage of variant sites = number of SNVs/(MAG length*breath)*). To characterize the distribution of strains and sharing events we used *compare* command from inStrain, which conducts paired comparisons of popANI metric (i.e., average nucleotide identity among locations along the genome where no alleles are shared between either sample). A popANI value ≥99.99% when MAG in both samples being compared had >5x and >50% breadth was considered evidence for strain sharing.

### Phylogenetic placement of MAGs and isolated strains

Contigs from each MAGs were used as queries in a blastX^63^ against the nr database from NCBI (version November 11/10/2023). Resulted best hits guided the choice of species reference genomes to retrieve from NCBI and include in the phylogenetic analysis, in addition to outgroups. Single-copy orthologs were recovered with OrthoFinder v.2.5.4^64^, aligned with MAFFT v.7.520^65^, trimmed with trimAl^66^ option *-automated1*, and then concatenated for IQ-TREE v.2.2.0.3^67^ maximum likelihood analysis, options *-m TEST -bb 1000.* The website iTol was used for tree visualization^68^. Similarly, phylogenies from Sanger sequenced amplicons of isolated strains (see below, “Bacteria and fungi isolation and taxonomy”) only skip the single-copy orthologs recovery step.

### Microbiome and MAG encoded functions

Contigs from each sample had coding sequences (CDS) predicted with Prodigal trough Prokka v.1.14.6^69^, using default options. Diamond v2.1.7.16^70^ was then used to blast the CDS against the nr database from NCBI (version November 11/10/2023), using -*-max-target-seqs 1 flag*, and the corresponding best hit taxa was recovered with NCBI Entrez API. The broad consensus taxa of each contig (*i.e.,* bacteria, fungi, insecta or other) was determined as the majority of contigs’ CDS taxa classification. Having a list of all bacterial contigs, we extracted their CDS sequences to run eggnog-mapper^71^, using emapper v.2.1.1 and eggNOG DB v.5.0.2, *-m diamond*. Predicted KEGG ortholog IDs (ko:K#####) for each protein were used to recover BRITE classification. To test for functional enrichment in each hive component, coordinates of each CDS were recovered from Prokka gff ouput file, and used to recover the mean coverage from contig depth file (see “Metagenome assembly and binning”) which was then normalized for reads per million (*RPM=mean coverage/total mean coverage)*1000000*). The CDS from MAGs were also predicted and imported to KBase^72^ to run DRAM^73^ and recover additional information about carbon metabolism.

### Bacteria and fungi isolation and taxonomy

Bacteria and fungi were isolated from larvae and larval diet samples using a variety of media recipes, with the final type used for each strain stated (**Supplementary Table 7**). Briefly, raw, macerated and PBS 1X diluted samples were plated in Petri dishes, then incubated at derived-bee species brood temperature (i.e., 30°C for stingless bee isolates^31^ and 34°C for honey bee isolates^32^). Plates were checked every day for microbial growth, then colonies, spores or mycelia pieces replated until isolation. Stocks of cells for bacteria and yeast or spores for filamentous fungi were stored in cryotubes at −70◦C with 20% glycerol. Strain isolation and taxonomy was confirmed by sequencing 16S rRNA gene for bacteria (primers: 27F 5′- AGAGTTTGATCCTGGCTCAG-3′ and 1492R 5′-TACGGYTACCTTGTTACGACTT-3′^74^) and 18S rRNA gene (primers: NS1 5′-GTAGTCATATGCTTGTCTC-3′ and NS4 5′- CTTCCGTCAATTCCTTTAAG-3′) or ITS region (primers: ITS5 5’ GGAAGTAAAAGTCGTAACAAGG 3’ and ITS4 5’ TCCTCCGCTTATTGATATGC 3’)^75^ for fungi, observing single clean peaks in chromatograms. A maximum likelihood phylogeny was inferred for *Zygosaccharomyces* strains as described in “Phylogenetic placement of MAGs and isolated strains” section. For *Starmerella*, only the sequence alignment was performed, as bootstrap support was insufficient for a reliable phylogenetic reconstruction due to limited number of closely related sequences in public databases.

### Culturing bacteria and fungi for *in vivo* and *in vitro* experiments

Three media types were used for strain cultivation and experiments: 30G (30 g glucose, 3 g malt extract, 3 g yeast extract, and 100 mL deionized water^19^), PDA (Difco™ Potato Dextrose Agar, 213400), and MRS (BD Difco™ Dehydrated Culture Media: Lactobacilli MRS Agar, 288210). Single bacteria and yeast colonies from each strain were transferred to 5 mL broth and grown for the experiments at their respective isolation temperature for 24 h, shaking at 230 rpm. After 24 h grow all bacterial strains reached 10^8^ cells/mL. Yeasts were gently centrifuged for 8 min, at 6000 rpm, and the cell pellet resuspended in 500 µL of sterile water for OD600 measurement and normalization. Filamentous fungi were grown in PDA agar under respective temperature until spores were observed. Spores were collected with sterile water and gently use of cell scraper, then counted with the help of a hemocytometer for normalization.

### *In vivo* larvae infection assays

Honey bee larvae were exposed to high loads of fungi cells or spores to test for potential pathogenicity to bees. Schmehl protocol^76^ was followed for larvae rearing, which currently is the only protocol using artificial larval diet. This ensured that no diet-derived microbes were present in the experiment, with only microbes inherent to the larvae or intentionally offered. Briefly, at day 0, first-instar larvae were grafted and deposited into plastic queen cups in 48-well plates, which already contained 10 µL of diet A. Larval diet was prepared no more than 24 h in advance of each feeding, always UV-sterilized for 20 min. On day 3, which is the first day of diet C, we added to each cup an additional 2 µL of 10^8^ fungal cells or spores/mL diluted in sterile water. For the negative control 2 µL of sterile water was added. At day 7, when larvae had finished to consume the last diet offer, they were transferred to a defecation 48-well plate with filter papers instead of queen cups. Development/survival was then followed until pupae reached brown-eyes stage, then weighed.

### Bacterial *vs* fungi competition assay

The experiment was conducted in 6-well agar plates, including two experimental conditions, as well as bacterial and fungal growth controls, each with three replicates. One experimental condition included a higher bacterial load (15 µL lawn of 10^8^ CFU/µL), while the other contained a lower bacterial load (15 µL lawn of 10^6^ CFU/µL). After the bacterial lawn had dried, a 5 µL drop containing yeast at OD600 = 0.5 or 500 spores was placed at the center of the well. The bacterial control received 5 µL of sterile water at the center, while the fungal control plate contained a lawn of MRS broth, the medium used for bacterial growth and dilutions. The experiment was incubated at 30°C for stingless bee bacterial strains and at 34°C for honey bee strains. Plates were monitored until the fungal positive controls show complete yeast growth or spores from filamentous fungi. Since *Zygosacharomyces* from stingless bees can only grow in 30G media, agar plates contained this media instead of MRS agar as used for the other fungi.

## Data availability

Raw metagenomic data has been deposited to the NCBI Sequence Read Archive (SRA) under the Project ID PRJNA1216660.

## Code availability

Custom scripts for our analysis are stored on GitHub https://github.com/liliancaesarbio/scaptotrigona_microbiome

## Acknowledgements

We thank beekeepers Evald Gossler, Nilo Techio, and Alcides Roque for their assistance with sampling from managed stingless bee hives, and Irmgard I. W. Caesar for providing hive photographs. We thank to the Laboratory of Molecular Biology at UNISINOS for technical support and to the Newton Lab members at Indiana University for their valuable feedback throughout this study. We also thank L. Felipe Benites, Chris Robinson and Carrie Ganote for insightful discussions on data analyses and writing. Finally, we acknowledge funding support: LC and CB were supported by the National Science Foundation (NSF) IOS Award #2005306 and the GEMS Biology Integration Institute, funded by the NSF DBI Biology Integration Institutes Program, Award #2022049. VHV was supported by the Conselho Nacional de Desenvolvimento Científico e Tecnológico (CNPq) (number 315532/2023-8).

## Author contributions

Conceptualization – LC, IN; Methodology – LC, IN; Experiments and Data Collection – LC, CB; Data Analysis – LC; Writing Original Draft – LC; Review and Editing – CB, VHV, IN; Resources – VHV, IN; Funding acquisition – LC, IN.

