## Supplementary Materials S1-S6 for "Spatial segregation and cross-kingdom interactions drive stingless bee hive microbiome assembly"

**Supplementary Figures**

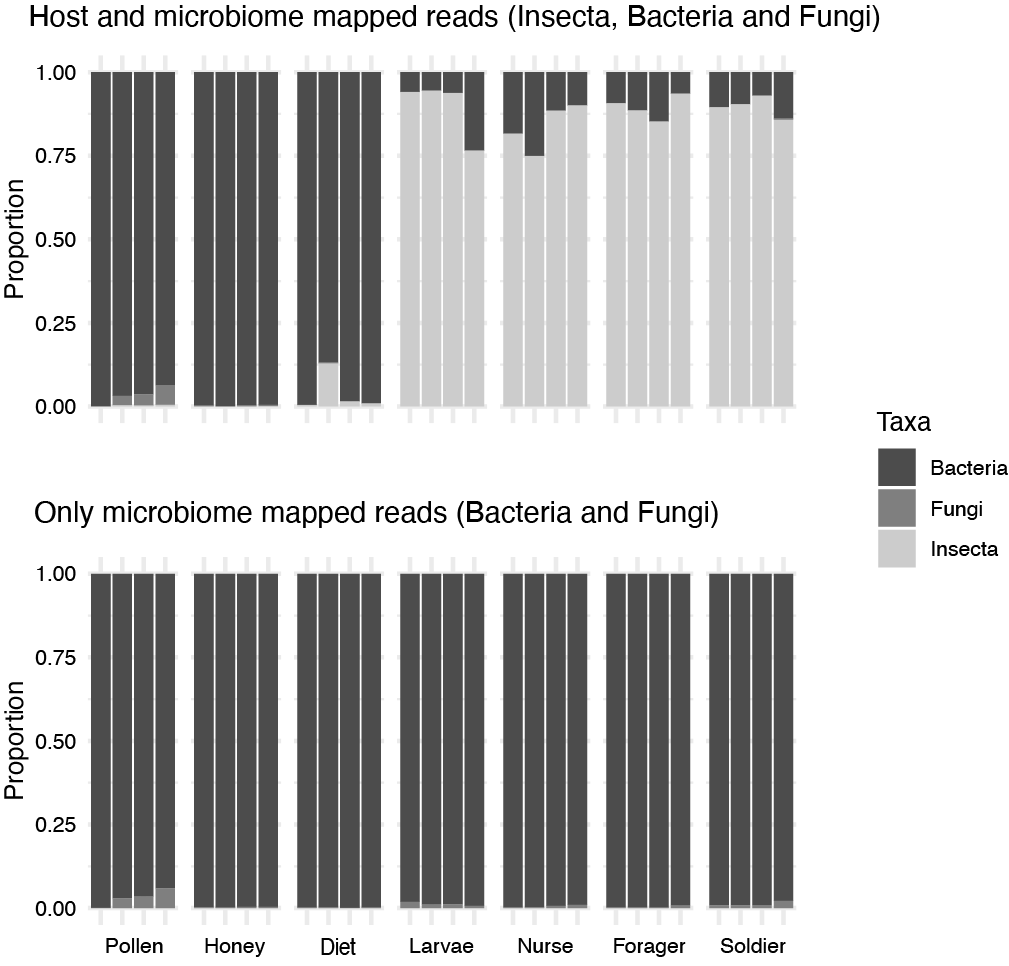

**Supplementary Fig. 1:** Proportion of reads mapping to Bacteria, Fungi, or Insecta from the SILVA SSU database v.138.1. As expected, pollen and honey samples show no Insecta reads, while some larval diet samples contain a small portion due to eggs laid in the brood comb. Bee samples have a significantly higher proportion of bee reads, as they include whole larvae and adult bee guts.

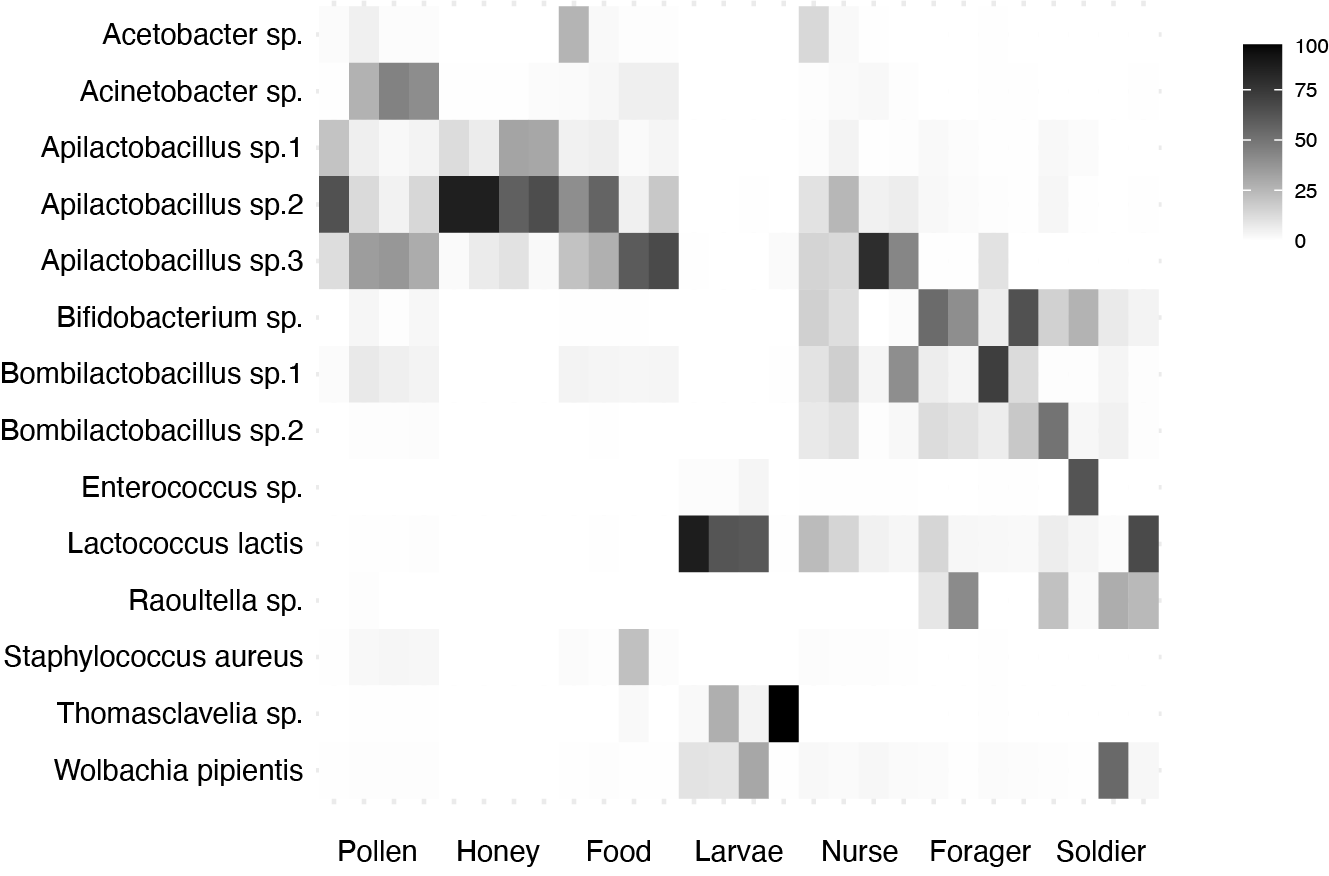

**Supplementary Fig. 2:** Proportion of all high-quality MAGs recovered in the 28 metagenome samples, including non-persistent strains. No bacteria associated with larvae exhibit coverage >1× or breadth >50%.

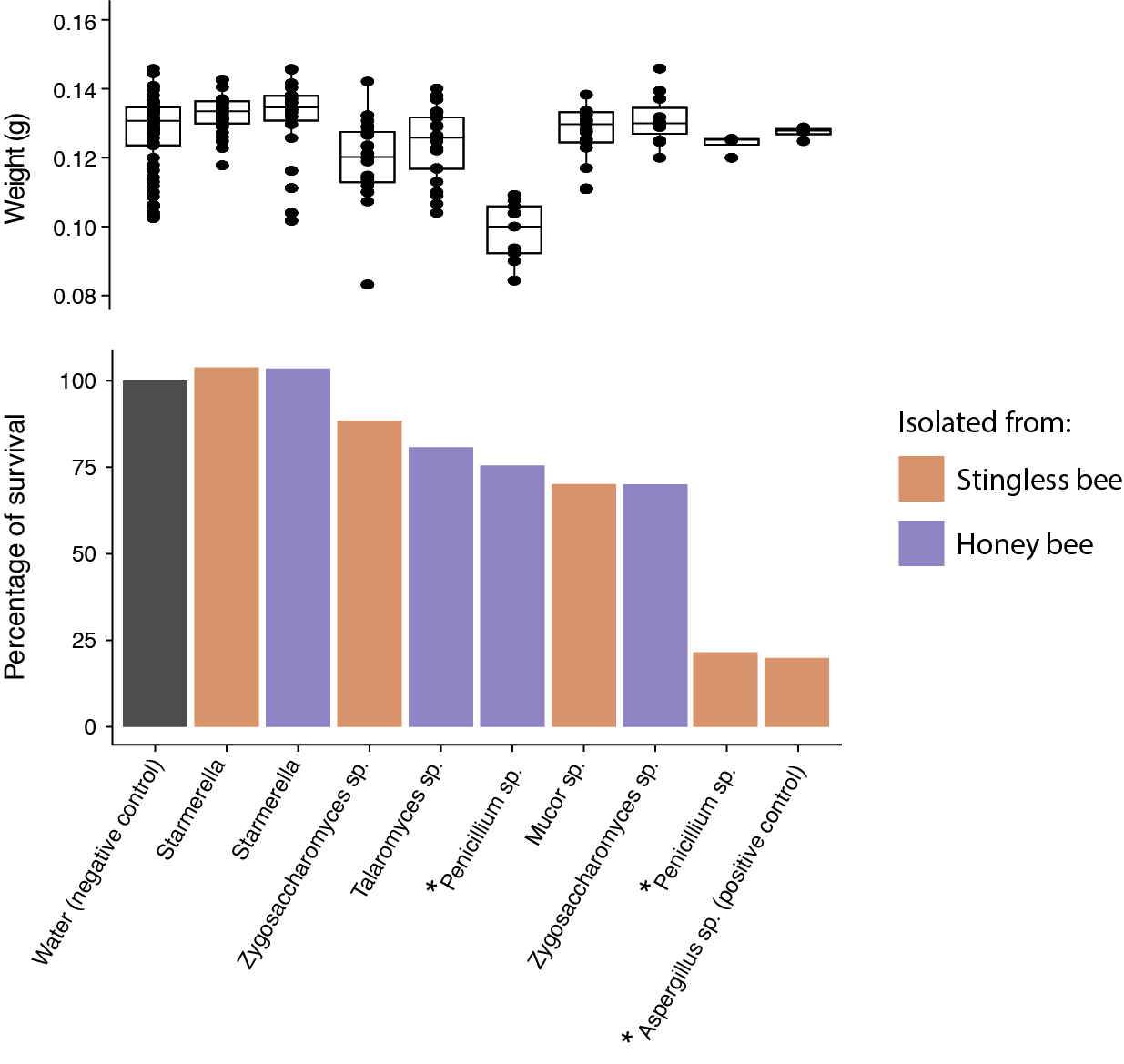

**Supplementary Fig. 3:** Larvae survival assay following challenge with a high load of fungal spores or yeast cells. Brown-eyed pupae that survived were weighed (above) and counted for survival percentage (below). Asterisks (*) indicate fungi potentially pathogenic to bees according to these results.

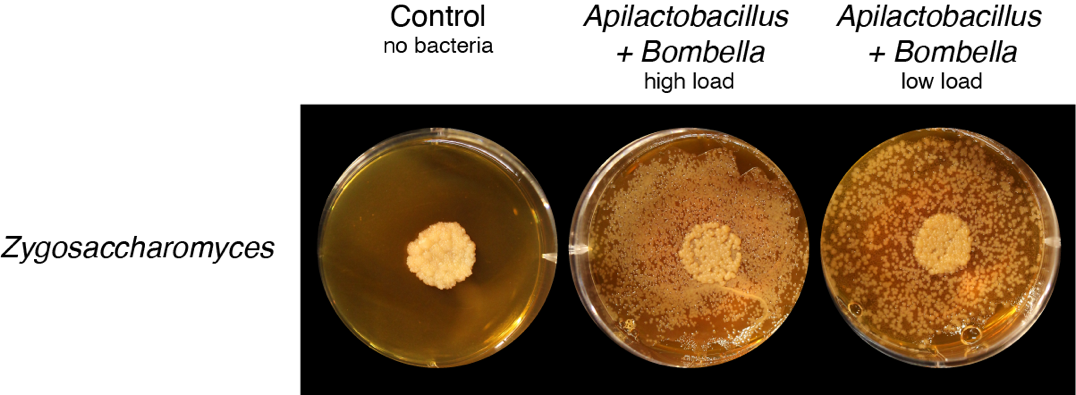

**Supplementary Fig. 4:** Competition assay between the *S. depilis* mock microbial community, composed of equal parts *Apilactobacillus* and Bombella (lawn), and the *Zygosaccharomyces* yeast symbiont (center). High (10⁸ CFU) and low (10⁶ CFU) bacterial loads were tested, and *Zygosaccharomyces* grew in both conditions.

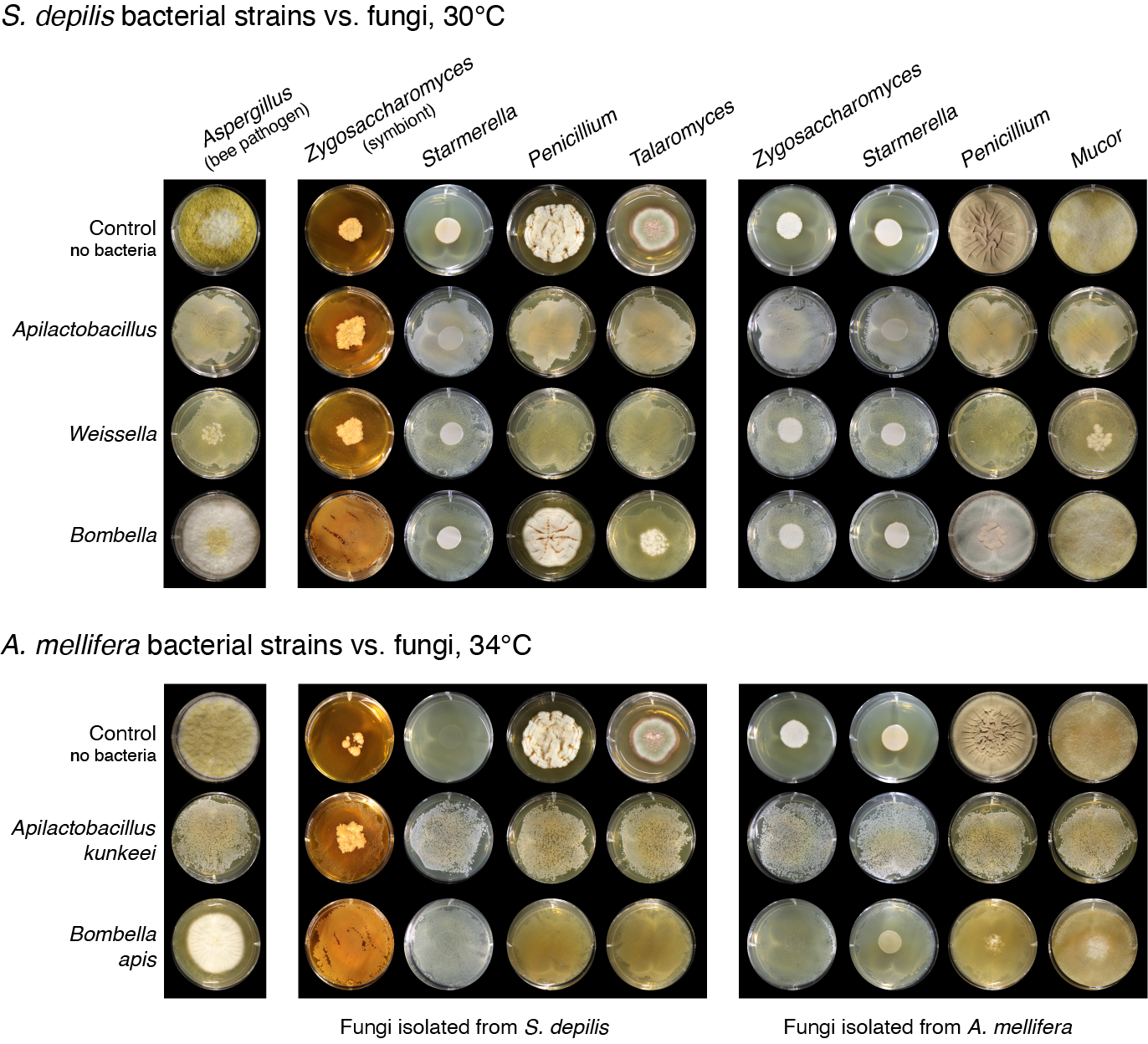

**Supplementary Fig. 5:** Competition assays between bacterial isolates from *Scaptotrigona depilis* larval diet (above) and *Apis mellifera* larvae (below) vs. fungal isolates, as shown in Figure 4 of the main manuscript. Here, however, the bacterial lawns have lower load; from 10⁶ CFU/mL.

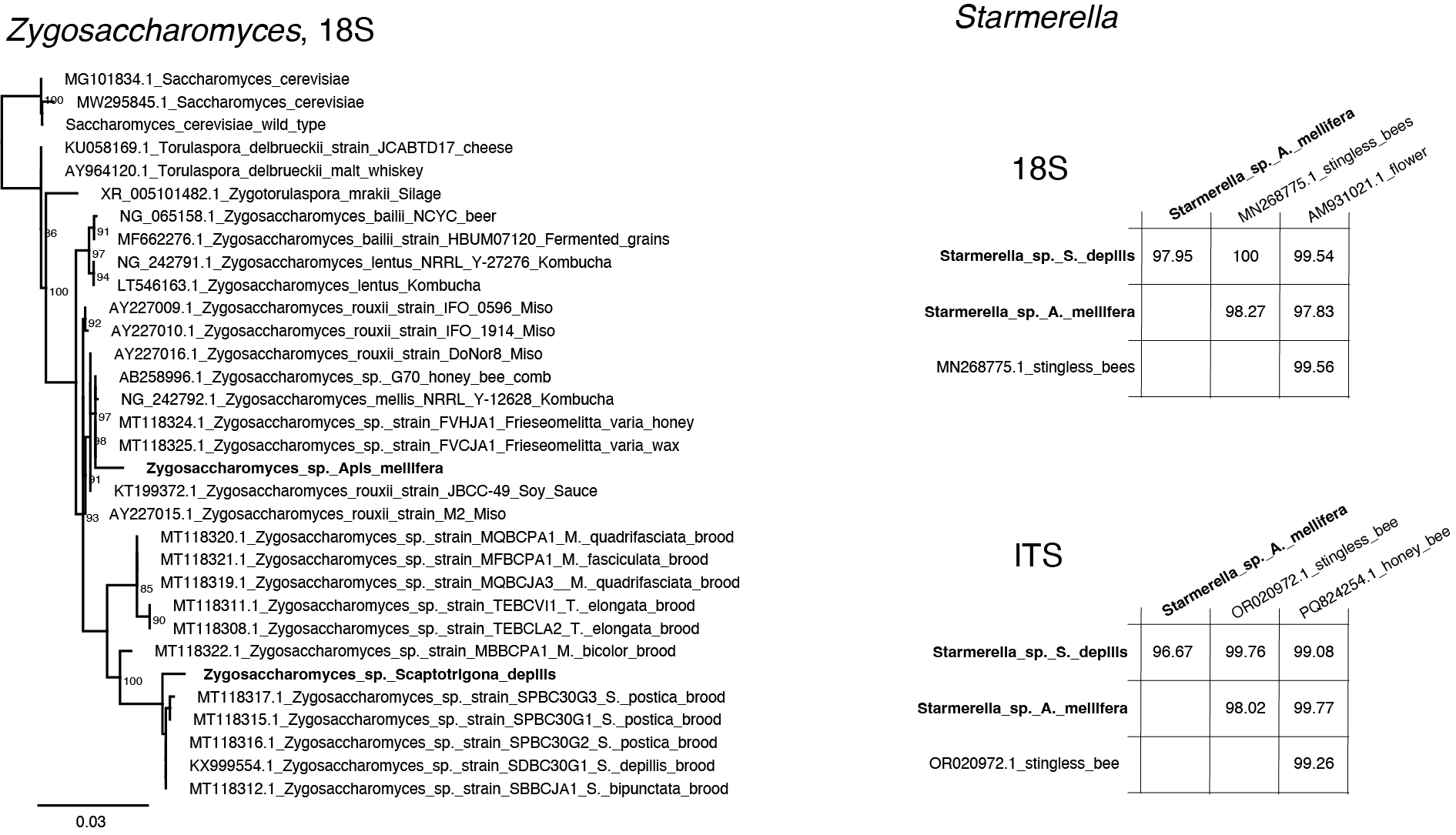

**Supplementary Fig. 6:** **On the left,** maximum likelihood phylogenetic analysis of *Zygosaccharomyces* based on the alignment of the 18S region. Bootstrap values >80 are indicated, and tree leaves contain sequence information followed by the isolation source. For *Zygosaccharomyces*, considerable effort has been made to sample strains associated with different stingless bee hive components, as well as other sources (e.g., food). In contrast, for *Starmerella*, there are only few studies and sequences available. The phylogenetic inference using ITS or 18S did not yield topologies with good bootstrap support, thus, **on the right**, we show a matrix with sequence identity to their closest strains after alignment and trimming out non informative sites. Sequence name contains ID number and isolation source. **Left and right**; sequences we isolated and used in this study are highlighted in bold.
